# Asparagine deprivation causes a reversible inhibition of Human Cytomegalovirus acute virus replication

**DOI:** 10.1101/690016

**Authors:** Chen-Hsuin Lee, Samantha Griffiths, Paul Digard, Nhan T. Pham, Manfred Auer, Juergen Haas, Finn Grey

## Abstract

As obligate intracellular pathogens, viruses rely on the host cell machinery to replicate efficiently, with the host metabolism extensively manipulated for this purpose. High throughput siRNA screens provide a systematic approach for the identification of novel host-virus interactions. Here, we report a large-scale screen for host factors important for human cytomegalovirus (HCMV), consisting of 6,881 siRNAs. We identified 47 proviral factors and 68 antiviral factors involved in a wide range of cellular processes including the mediator complex, proteasome function and mRNA splicing. Focused characterisation of one of the hits, asparagine synthetase (ASNS), demonstrated a strict requirement for asparagine for HCMV replication which leads to an early block in virus replication before the onset of DNA amplification. This effect is specific to HCMV, as knockdown of ASNS had little effect on herpes simplex virus-1 or influenza A virus replication, suggesting the restriction is not simply due to a failure in protein production. Remarkably, virus replication could be completely rescued seven days post-infection with addition of exogenous asparagine, indicating that while virus replication is restricted at an early stage, it maintains the capacity for full replication days after initial infection. This study represents the most comprehensive siRNA screen for the identification of host factors involved in HCMV replication and identifies the non-essential amino acid, asparagine as a critical factor in regulating HCMV virus replication. These results have implications for control of viral latency and the clinical treatment of HCMV in patients.

**Importance:** HCMV accounts for more than 60% of complications associated with solid organ transplant patients. Prophylactic or preventative treatment with antivirals, such as ganciclovir, reduces the occurrence of early onset HCMV disease. However, late onset disease remains a significant problem and prolonged treatment, especially in patients with suppressed immune systems, greatly increases the risk of antiviral resistance. Very few antivirals have been developed for use against HCMV since the licensing of ganciclovir, and of these, the same viral genes are often targeted, reducing the usefulness of these drugs against resistant strains. An alternative approach is to target host genes essential for virus replication. Here we demonstrate that HCMV replication is highly dependent on levels of the amino acid asparagine and knockdown of a critical enzyme involved in asparagine synthesis results in severe attenuation of virus replication. These results suggest that reducing asparagine levels through dietary restriction or chemotherapeutic treatment could limit HCMV replication in patients.

## Introduction

Human cytomegalovirus (HCMV) is a highly prevalent herpesvirus, infecting greater than 30% of the worldwide population. Although normally asymptomatic in healthy individuals, HCMV infection is a significant cause of morbidity and mortality in immunocompromised populations, individuals with heart disease and recipients of solid organ and bone marrow transplants (1). HCMV is also the leading cause of infectious congenital birth defects resulting from spread of the virus to neonates (2).

Cellular metabolism is a tightly regulated process in mammalian cells and is often manipulated during viral infection. As obligate intracellular pathogens, viruses rely on host metabolites and often alter host metabolism to increase pools of free nucleotides and amino acids, as well as inducing fatty acid biosynthesis to aid efficient virus replication (3, 4).

Infection with HCMV has been demonstrated to alter the host cell metabolic pathways, increasing glycolysis and glutamine metabolism while maintaining protein translation through activation of mammalian target of rapamycin complex 1 (mTORC1) (5). Glucose metabolism is a key pathway to supply carbon precursors for cellular biosynthesis and energy production. Under normal conditions, glucose is used for energy generation and cellular biosynthesis whilst only a small amount of glutamine is metabolised from exogenous sources. In contrast, in HCMV infected cells, glucose is diverted away from TCA cycle, into the production of lactic acid and fatty acid, while exogenous glutamine is used as the main carbon and nitrogen source. Glutamine can donate its amino group at the gamma position for *de novo* biosynthesis of nucleotides and non-essential amino acids whilst being converted to glutamate (6). Glutamate can be further metabolised into α-ketoglutarate via glutamate dehydrogenase, thereby providing a key intermediate for the TCA cycle, a process known as anaplerosis, which also occurs in rapidly dividing cancer cells (7).

A recent study showed that infection with HCMV results in increased metabolism of arginine, leucine/isoleucine, serine and valine and increased secretion of alanine, ornithine and proline, demonstrating extensive alteration of cellular amino acid metabolism during infection (8). Furthermore, HCMV manipulates cellular signalling pathways to maintain protein synthesis during amino acid starvation. Mammalian cells have two main pathways that monitor and modulate the level of intracellular amino acids: mTOR and the amino acid response (AAR) pathway. The mTOR pathway serves to ensure a sufficient level of amino acids to support protein synthesis and cell growth. Previous studies have shown that glutamine and leucine activate the mTOR pathway via glutaminolysis and mediate cellular responses to amino acids (9). Activation of mTOR ultimately leads to the phosphorylation and activation of the ribosome-associated S6 kinase, which enables higher levels of protein synthesis, while loss of mTOR signalling results in suppression of protein synthesis. However, HCMV infection can maintain mTOR activation during amino acid deprivation, through the viral UL38 protein binding and antagonising the tuberous sclerosis subunit complex 2 (TSC2), a major suppressor of mTOR (10). UL38 interaction with TSC2 has also been shown to have broader effects on cellular metabolism in an mTOR independent fashion (8). These findings show that regulation of amino acid metabolism plays an important role during HCMV replication.

Here, we show that asparagine synthetase (ASNS) is a critical host factor for HCMV replication following a comprehensive siRNA screen. Knockdown of ASNS resulted in an early restriction in virus replication. However, knockdown of ASNS had little effect on Herpes Simplex Virus-1 (HSV-1) or influenza A virus (IAV) replication, indicating the effects of asparagine depletion was specific to HCMV and not simply due to a loss of production of asparagine-containing proteins. Furthermore, mTOR activation was maintained in infected cells following ASNS knockdown, indicating that this was not the cause of attenuated virus replication. Remarkably, the block in viral replication could be completely rescued seven days post-infection with the addition of exogenous asparagine to the cell media. These results suggest a novel check point in virus replication regulated by intracellular asparagine levels.

## Results

### High-throughput siRNA screen identified novel host factors involved in HCMV replication

To identify host factors that influence HCMV replication, a combined siRNA library comprising small interfering RNAs (siRNAs) targeting the human druggable genome, protein kinases/phosphatases and cell cycle genes, were used in a high-throughput screen. SMARTpool siRNAs (a pool of 4 siRNAs per gene) targeting a total of 6,881 genes were transfected into primary normal human dermal fibroblast (NHDF) cells in a 384 well format. Cells were infected at 48 hours post-transfection at a high multiplicity of infection (MOI = 5) with the low-passage HCMV strain TB40/E-GFP, which expresses green fluorescent protein (GFP) from a simian virus 40 (SV40) promoter (11). GFP fluorescence levels were monitored every 24 hours for seven days, with levels compared to control non-targeting siRNA transfected cells in order to determine the effect of individual gene depletion on HCMV replication. We have previously established GFP expression as an accurate measure of early virus replication events including viral entry, translocation to the nucleus and viral DNA amplification, hereon referred to as primary replication (12, 13). The assay was performed in biological triplicate with an additional replicate used to determine cytotoxicity at seven days post-infection (DPI) (supplemental table 1). siRNAs were defined as cytotoxic when gene depletion led to greater than 40% cell death (106 in total; data not shown). These genes were excluded from further analyses. High correlation between biological triplicates demonstrated reproducibility within the screen (**Figure 1A**). The screen identified a total of 115 host factors where knockdown led to 50% inhibition (47 proviral factors) or 50% increase (68 antiviral factors) in primary replication (**Figure 1B and C, supplemental table 2 and 3**). Figure 2 shows a schematic summary of the top hits affecting HCMV virus replication grouped into functionally related gene clusters based on STRING analysis (14). These include members of the proteosome complex and ubiquitin modifying factors, which have previously been shown to be important for efficient HCMV replication and the mediator complex that has been linked to immediate early transactivator function in alpha and gamma herpesviruses (15-19). mRNA splicing factors, transcription and translation initiation factors and DNA replication factors are also identified as proviral. Interestingly, a number of histone modification factors are identified as antiviral, along with multiple signal transduction factors, developmental, cell adhesion and cellular response to signalling factors. Based on the magnitude of phenotype, lack of toxicity and novelty, nine host factors with proviral phenotypes were selected for further validation (**Table 1**).

**Table 1.** siRNA deconvolution validation of nine proviral candidates. Nine proviral candidates were selected for validation with deconvoluted siRNAs. 4 individual siRNAs targeting different regions of each gene were used to transfect NHDF cells, followed by infection 48 hours post-transfection with HCMV TB40/E-GFP at an MOI of 5. The number of deconvoluted siRNAs that showed the same phenotype as the pool is shown. Individual siRNA was considered validated when knockdown led to significant inhibition of virus replication. Std. Dev. = standard deviation; *n* = 3

| Gene symbol | Mean Ratio to siNeg |  | Std. Dev. |  | siRNA deconvolution (out of 4) | Description |
| --- | --- | --- | --- | --- | --- | --- |
|  | Screen pool | Validation pool | Screen pool | Validation pool |  |  |
| SON | 0.14 | 0.09 | 0.10 | 0.02 | 4 | SON DNA binding protein |
| CSE1L | 0.15 | 0.16 | 0.03 | 0.02 | 3 | CSE1 chromosome segregation 1-like (yeast) |
| SNU13 | 0.20 | 0.11 | 0.09 | 0.01 | 4 | NHP2 non-histone chromosome protein 2-like 1 (S. cerevisiae) |
| GPS1 | 0.36 | 0.40 | 0.15 | 0.04 | 1 | G protein pathway suppressor 1 |
| UBA52 | 0.36 | 0.48 | 0.06 | 0.08 | 3 | Ubiquitin A-52 residue ribosomal protein fusion product 1 |
| ERH | 0.38 | 0.45 | 0.01 | 0.05 | 3 | Enhancer of rudimentary homolog (Drosophila) |
| PABPN1 | 0.46 | 0.50 | 0.10 | 0.08 | 3 | Poly(A) binding protein, nuclear 1 |
| ASNS | 0.47 | 0.18 | 0.22 | 0.02 | 3 | Asparagine synthetase |
| SEM1 | 0.49 | 0.64 | 0.03 | 0.04 | 4 | Split hand/foot malformation (ectrodactyly) type 1 |

**Figure 1.**
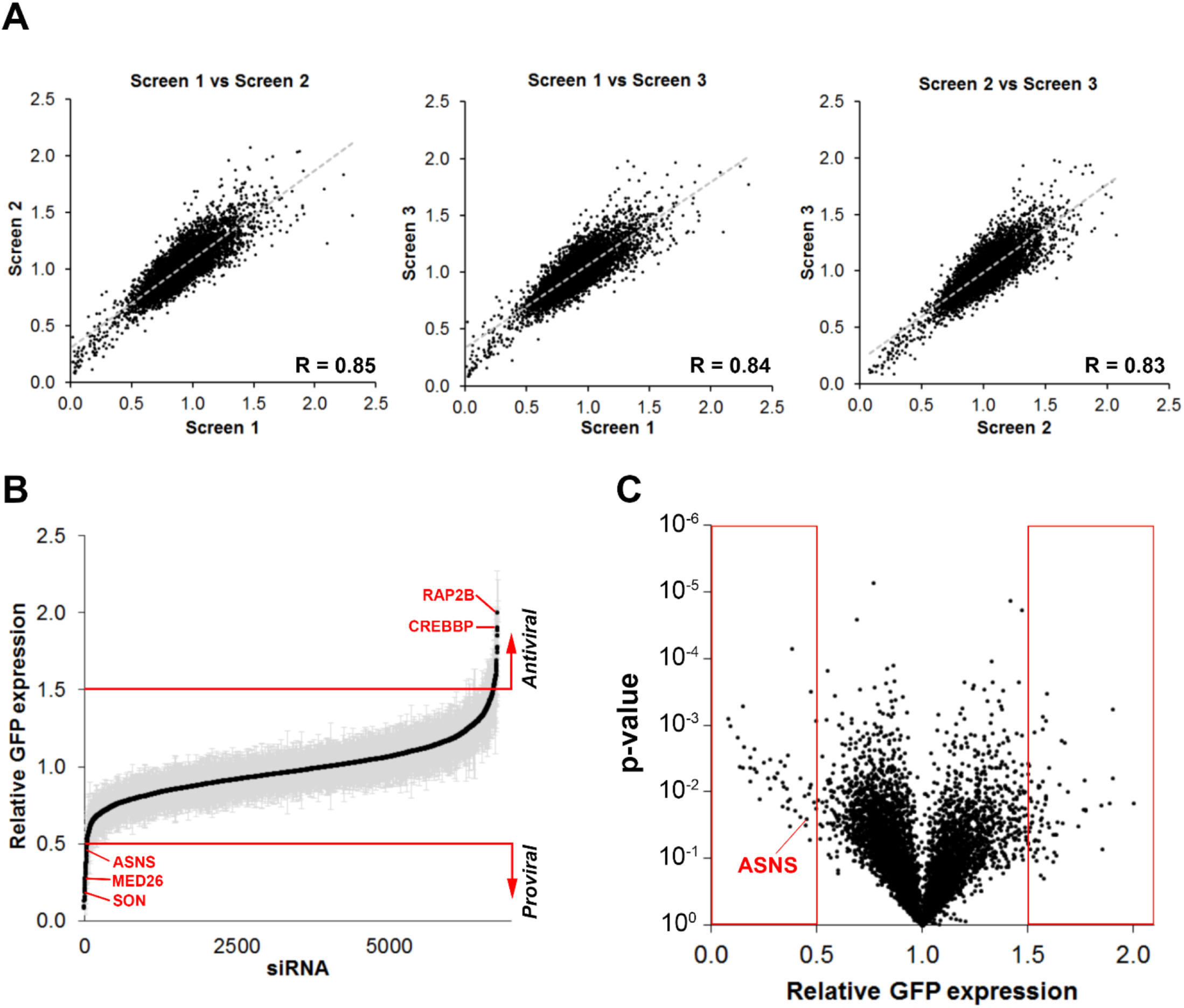
High-throughput siRNA screen identified novel host factors for HCMV replication. (A) NHDF cells were transfected with siRNA pools targeting 6,881 genes in a 384 well format.. Two days post transfection the cells were infected with HCMV TB40/E-GFP at an MOI of five with GFP levels monitored by plate cytometry for seven days. Comparative analysis between three biological repeats showed high levels of correlation with Pearson coefficient scores between 0.83 – 0.85. (B) Relative GFP expression representing the level of primary replication is shown, sorted from low to high compared to control non-targeting siRNA transfected cells. Standard deviations are shown in grey error bars. (C) Volcano plot showing relative GFP expression versus associated P-value. Each dot represents knockdown of a single gene. P-values were calculated by student’s *t*-tests. Red boxes represent the top hits based on a two-fold increase or decrease in relative GFP expression (listed in supplemental table 2 and 3).

**Figure 2.**
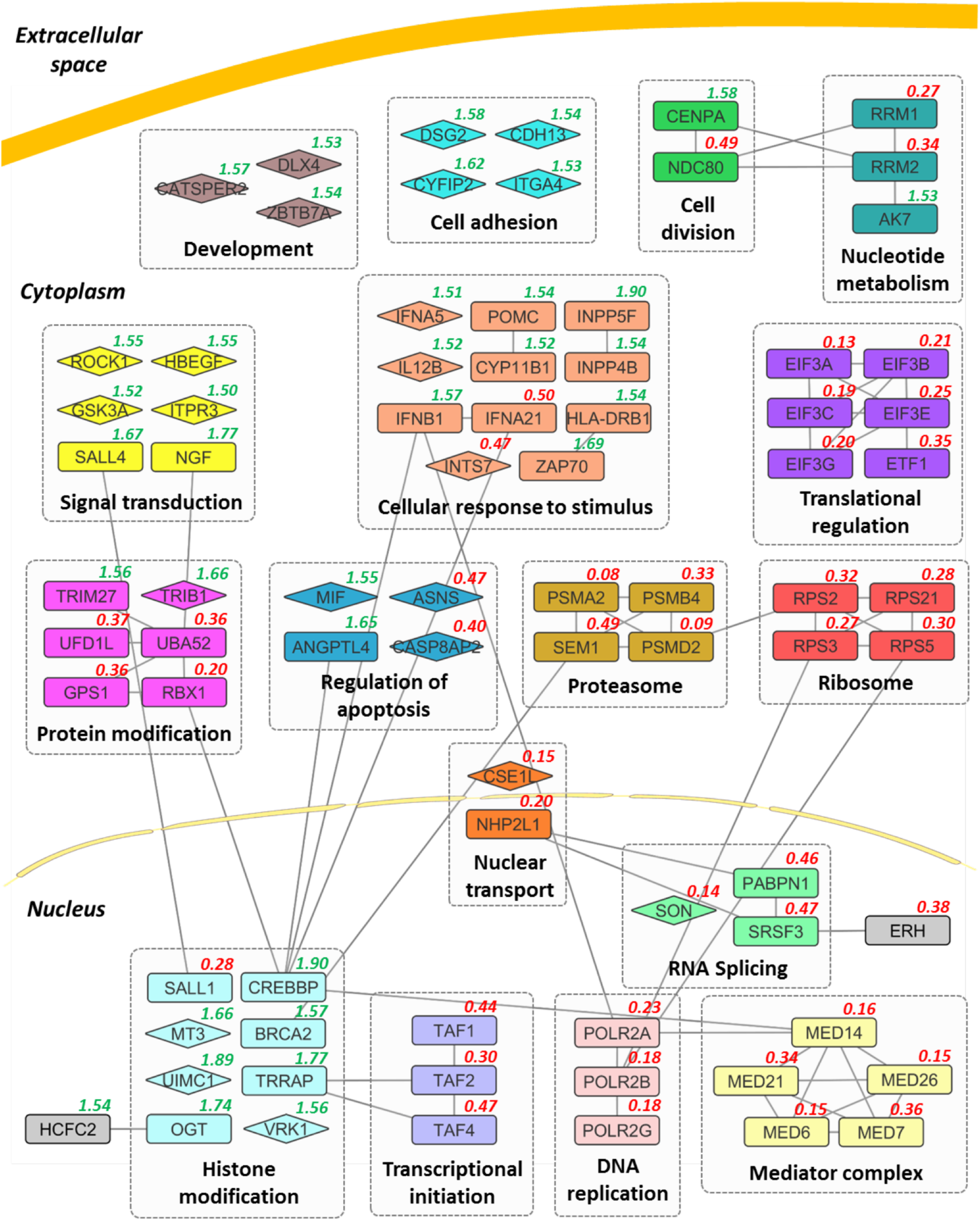
Functional analyses of top proviral and antiviral candidates. Selected candidates of the top 115 host factors are shown in functional clusters based on STRING analysis. Interactions between genes all have high interaction score (combined score >0.8, STRING). Genes that do not show an interaction with other genes in this figure are depicted as diamonds. Functional gene clusters are shown in their approximate cellular location with the primary replication ratio compared to negative control indicated (green indicates antiviral role, red indicates proviral role).

To determine whether the observed effects on virus replication were specific to the knockdown of the identified gene and not due to artefactual off-target effects, the pools of siRNAs were deconvoluted and each of the individual siRNAs tested for their effect on virus replication, along with the original pool of siRNAs (**Table 1**). All pooled siRNAs recapitulated the phenotype in the primary screen, once again confirming reproducibility of the assay. In eight out of nine cases, at least 3 of 4 siRNAs from the original pools demonstrated similar phenotypic effects, strongly suggesting attenuation of primary replication was due to knockdown of the target host factor rather than off-target effects. Asparagine synthetase (ASNS) was selected for further detailed characterisation as knockdown resulted in a substantial reduction in primary replication based on GFP expression, and has not previously been associated with HCMV or herpesviruses in general.

### ASNS is a crucial host factor for HCMV replication

ASNS is the sole enzyme that catalyses the biosynthesis of asparagine, by transferring the gamma amino group from glutamine to aspartate in an ATP-dependent manner. In the high throughput screen, knockdown of ASNS inhibited HCMV primary replication, based on GFP expression, throughout the course of a single cycle infection (**Figure 3A**). Inhibition was not due to siRNA cytotoxicity, as measured by CellTiter-Blue cytotoxicity assay (**Figure 3B**), with cell viability increased following ASNS knockdown in infected cells compared to control transfected cells, possibly due to the inhibition of HCMV replication and subsequent decrease in cytopathic effect in infected cells.

**Figure 3.**
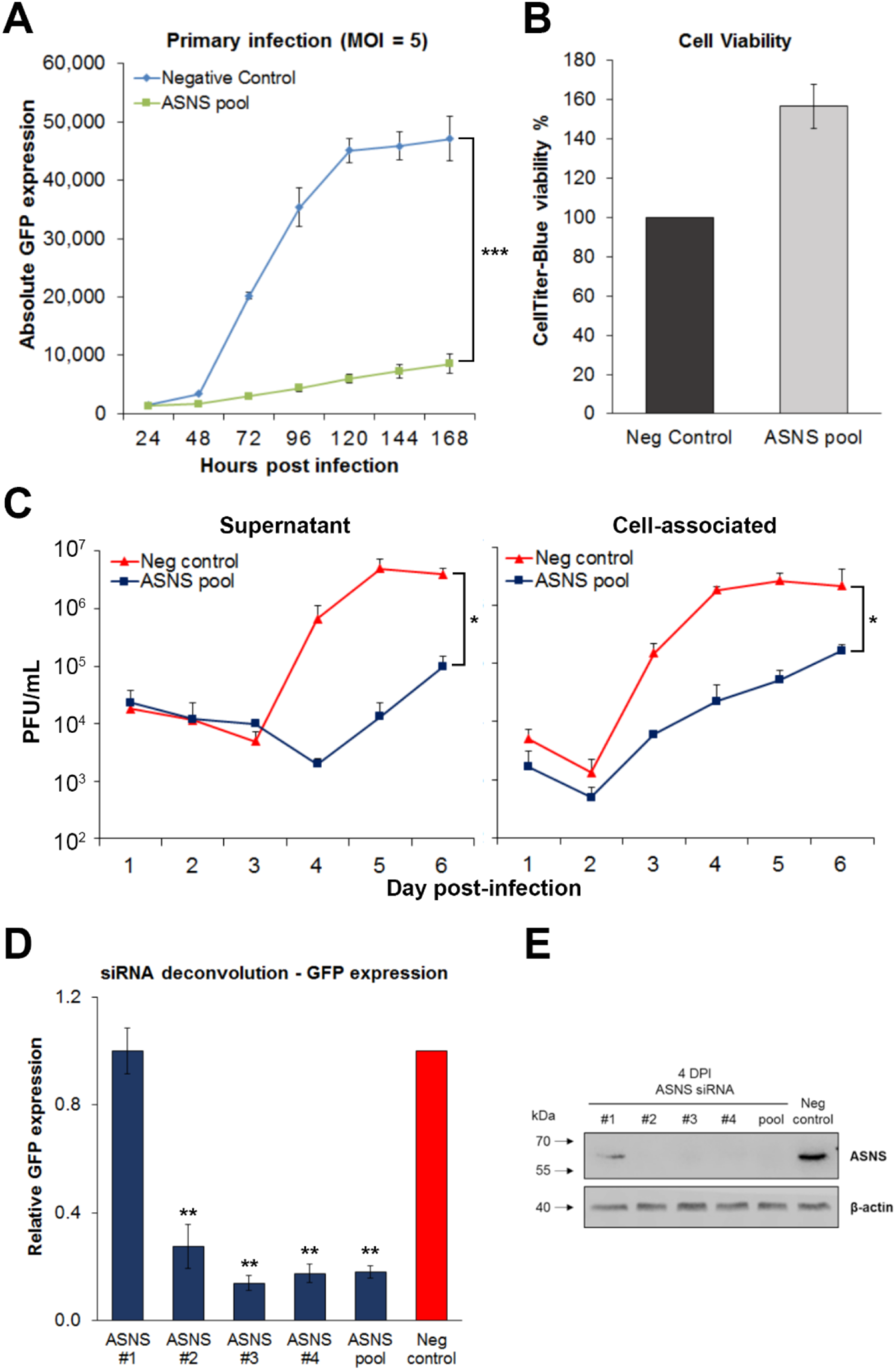
Asparagine synthetase (ASNS) is a novel host factor for HCMV replication. NHDF cells were transfected with an siRNA pool targeting ASNS or a scrambled negative control and infected 48 hours post-transfection with HCMV TB40/E-GFP at an MOI of 5. (A) Knockdown of ASNS substantially decreased virus replication throughout the course of infection. *n* = 3; error bars show standard deviations. Two-way ANOVA was used to calculate statistical significance. *** = p < 0.0005. (B) CellTiter-Blue assay was performed at 7 day post-infection (DPI) and cell viability was calculated. *n* = 3; error bars show standard deviations. (C) Cell-free (supernatant) and cell-associated virus (cells) were harvested at indicated time points, and virus levels determined by plaque assay. *n* = 2; error bars represent standard error of the mean. Two-way ANOVA was used to calculate statistical significance. * = p < 0.05. (D) NHDF cells were transfected with the 4 individual siRNAs targeting ASNS or a scrambled sequence (Neg control) and infected 48 hours post-transfection with TB40/E-GFP at an MOI of 5. GFP levels were measured at 7 DPI to determine the effect of gene depletion on primary replication. *n* = 3; error bars represent standard deviations. One-way ANOVA was used to calculate statistical significance between individual siRNA and the negative control. ** = p < 0.005. (E) Western blot analysis showed ASNS protein levels following knockdown with individual siRNA (1-4) and pool, along with negative control. Protein lysates were collected 4 DPI.

To confirm reduced GFP expression levels corresponded to reduced virus production, single-step growth curves were performed following knockdown of ASNS. Supernatant and cell-associated virus levels were determined following a high multiplicity infection (MOI = 5) by plaque assay. In both pools, knockdown of ASNS resulted in a substantial decrease in virus production compared to control non-targeting siRNA transfected cells, confirming the original observation based on GFP expression (**Figure 3C**).

In order to confirm the observed phenotype was due to specific protein knockdown and not due to off-target effects, we performed individual siRNA knockdown with 4 separate siRNAs from the pool. Three out of four siRNAs against ASNS showed the same phenotype as the reconstituted siRNA pool, whereas siRNA #1 showed no inhibition of primary replication (**Figure 3D**). Western blot analysis with protein lysates collected at 96 hours post-infection (HPI) revealed that siRNA #1 was less efficient at knocking down the protein, demonstrating a direct correlation between siRNA efficacy and attenuation of virus replication (**Figure 3E**), confirming that the observed phenotype was due to specific knockdown of ASNS and not off-target effects.

### ASNS knockdown inhibits HCMV IE2 and subsequent gene expression

To determine where in the virus lifecycle ASNS knockdown restricts virus replication, the associated phenotype was characterised in more detail. Knockdown of ASNS did not significantly affect viral entry or translocation of the genome to the nucleus as the number of GFP positive cells were equivalent between ASNS knockdown and control transfected cells at 24 hours post infection (**Figure 4A**). Thereafter, we investigated the role of ASNS in HCMV gene expression in more detail. HCMV gene expression occurs in a temporal manner, with immediate-early (IE), early (E) and late (L) gene expression phases. In order to identify the stage of the HCMV life cycle ASNS was involved in, we qualitatively measured the expression of IE (IE1/2), E (pp52) and L (pp28) genes by western blot analysis following knockdown of ASNS (**Figure 4B & 4C**). Following ASNS knockdown, IE1 expression was not drastically affected compared to the control non-targeting transfected cells. However, IE2 expression was substantially reduced, with subsequent E and L gene expression reduced to below the level of detection. Unsurprisingly, viral DNA levels were also significantly reduced following ASNS knockdown (**Figure 4D**). These results indicate that knockdown of ASNS results in an early phenotype with inhibition of virus replication occurring after viral entry and translocation to the nucleus, but before IE2 gene expression and viral DNA amplification.

**Figure 4.**
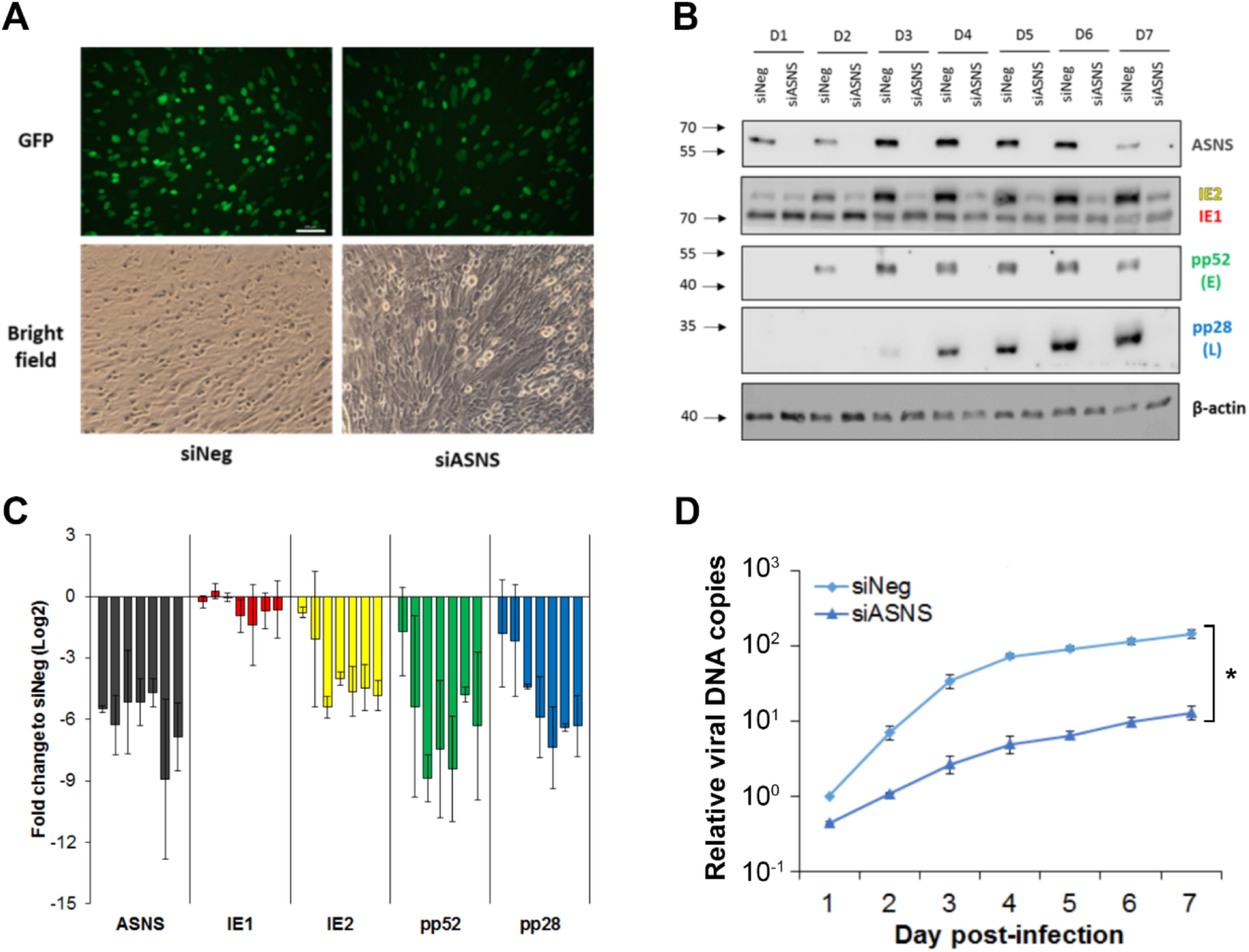
Knockdown of ASNS reduced IE2 expression and viral DNA amplification. NHDF cells were transfected with siRNA pool targeting ASNS (siASNS) or a scrambled negative control siRNA (siNeg), followed by infection 48 hours post-transfection with TB40/E-GFP at an MOI of 5. (A) Fluorescence images were taken at 1 DPI with 100X magnification. White bar = 100 µm. (B) Total protein was harvested at indicated time points and levels of immediate-early (IE1/2), early (pp52) and late (pp28) proteins were detected by western blot analysis. (C) Quantification of western blot data from figure B. The relative intensity of each band was normalised to β-actin compared to siNeg at day 1. Bars in each gene represent the differential expressions from 1 to 7 DPI (left to right). *n* = 2 (D) Total genomic DNA was isolated at indicated time points, and viral genome levels were determined by qPCR. *n* = 2; error bars show standard errors of mean. Two-way ANOVA was used to calculate statistical significance. * = p < 0.05.

### Inhibition of HCMV replication is not due to a general loss of protein translation

As standard growth media does not contain asparagine, NHDF cells are dependent on ASNS for generation of *de novo* asparagine. Knockdown of ASNS would therefore be predicted to lead to asparagine depletion over time, ultimately impacting translation of proteins containing asparagine. However, the relatively early phenotype observed and the fact that IE1 protein expression levels were not dramatically affected, while IE2 expression levels were substantially reduced, suggests that the effect on HCMV replication following ASNS knockdown may not be due to a general loss of protein translation caused by asparagine deprivation. To directly measure protein translation levels in infected cells following ASNS knockdown, puromycin pulse studies were performed. Cells were transfected with siRNA against ASNS or a negative control siRNA and infected with TB40E-GFP at an MOI of five, two days post transfection. Puromycin was added at zero (time of infection), four and seven days post infection (DPI) for fifteen minutes, before total proteins were harvested. Global translation levels were measured by western blot analysis using a puromycin specific antibody. In addition to general labelling of proteins, a strong band was detected at approximately 24 kDa that is substantially reduced in the ASNS knockdown cells (**Figure 5A**). Currently the identity and relevance of this protein is not known, however experiments are ongoing to identify the protein and determine whether it plays a direct role in ASNS dependant HCMV replication. To avoid skewing the analysis, this band was omitted from the quantification analysis. Other than this band, the results show that relative protein translation levels were not reduced following ASNS knockdown in infected cells (**Figure 5A and 5B**). This suggests that inhibition of HCMV replication from ASNS knockdown is not simply due to reduced protein translation due to the loss of available asparagine, but rather the virus is responding in a more indirect way to asparagine levels within the cell. Furthermore, knockdown of ASNS does not reduce replication of the related herpes simplex virus-1 (HSV-1) or influenza A virus (IAV). Following knockdown of ASNS, primary human fibroblast cells were infected with HSV-1 (strain C12) or IAV (strain A/PR/8/34 [PR8]) at 48 hours post-transfection (**Figure 5C and D**). In contrast to HCMV, HSV-1 and IAV replication were not significantly reduced, indicating the cells are still capable of supporting robust virus replication and the effect of ASNS knockdown is specific to HCMV.

**Figure 5.**
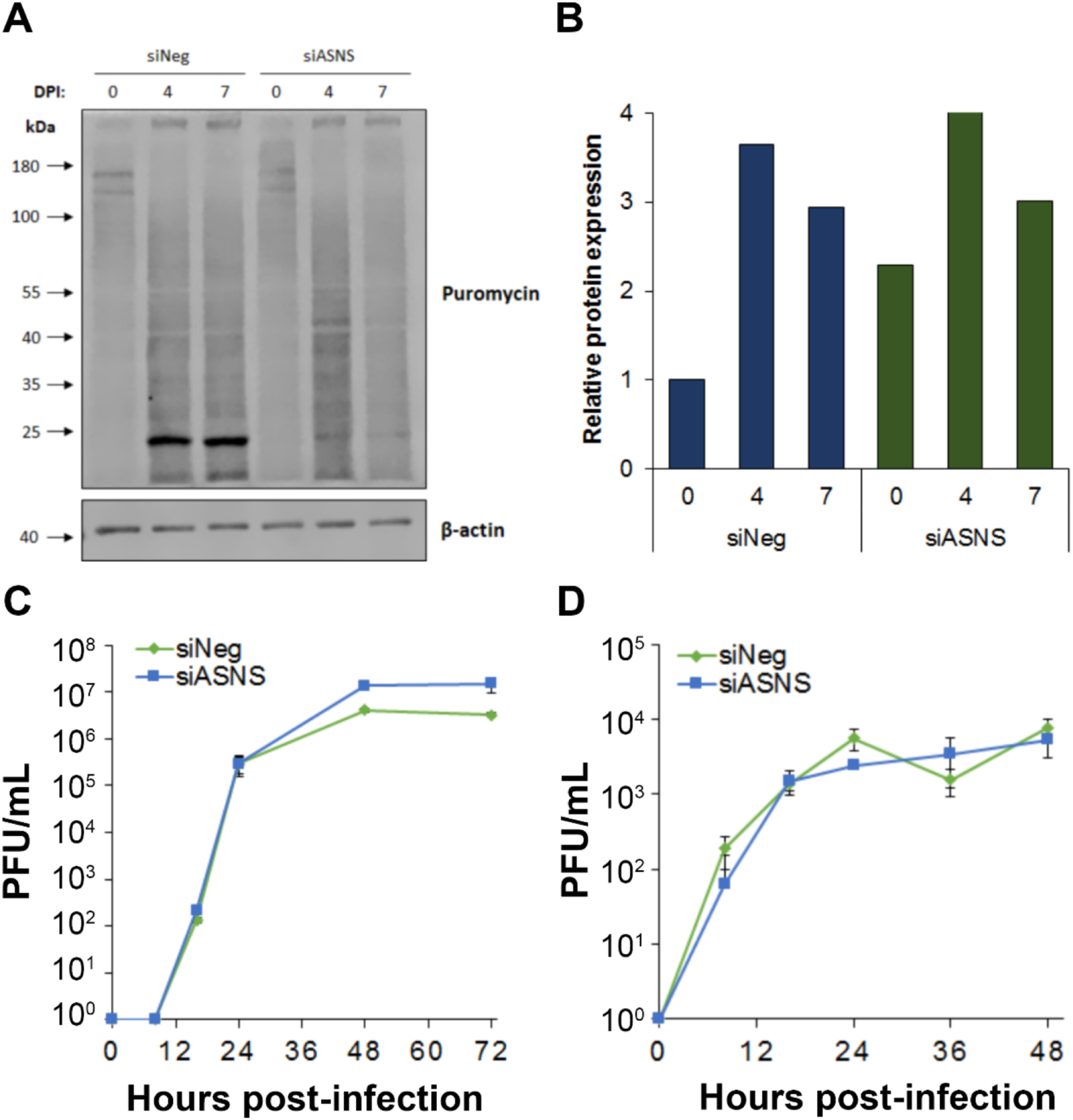
Inhibition of HCMV following ASNS knockdown is not due to loss of protein translation. (A) NHDF cells were transfected with siRNA pool targeting ASNS (siASNS) or a scrambled negative control siRNA (siNeg), then infected 48 hours post-transfection with HCMV TB40/E-GFP at an MOI of 5. Cells were treated with puromycin at indicated times. Global protein translation levels were quantified based on puromycin incorporation by western blot analysis (B) Quantification of western blot data from figure 5A. The relative intensity of each band was normalised to β-actin compared to siNeg at day 0. NHDF cells were transfected as above then infected with HSV-1 (C) or IAV (D) at an MOI of 0.1. Cells were harvested at the indicated times and virus levels quantified by plaque assay, n=3, error bars show standard deviation.

### Knockdown of ASNS did not inhibit mTOR activity during HCMV replication

As a reduction in protein translation does not appear to explain the inhibition of HCMV replication following ASNS knockdown, asparagine levels may lead to an indirect inhibition of HCMV virus replication through signalling pathways that monitor amino acid levels. While previous studies have shown that HCMV can override cellular signals that would normally inhibit mTOR activation due to deprivation of essential amino acids, the effect of deprivation of non-essential amino acids has not been investigated. Asparagine has recently been shown to function as an amino acid exchange factor and an indirect regulator of mTOR signalling (**Figure 6A**) (20). We therefore investigated mTOR signalling in ASNS depleted cells during HCMV replication to determine whether depletion of the non-essential amino acid asparagine had an inhibitory effect on its activation. mTOR signalling was maintained upon ASNS knockdown, based on phosphorylation of ribosomal S6 kinase (S6K) which is the downstream effector of the mTOR complex (**Figure 6B**). This suggests that the inhibitory effect of HCMV primary replication caused by ASNS depletion was not due to a loss of mTOR activation.

**Figure 6.**
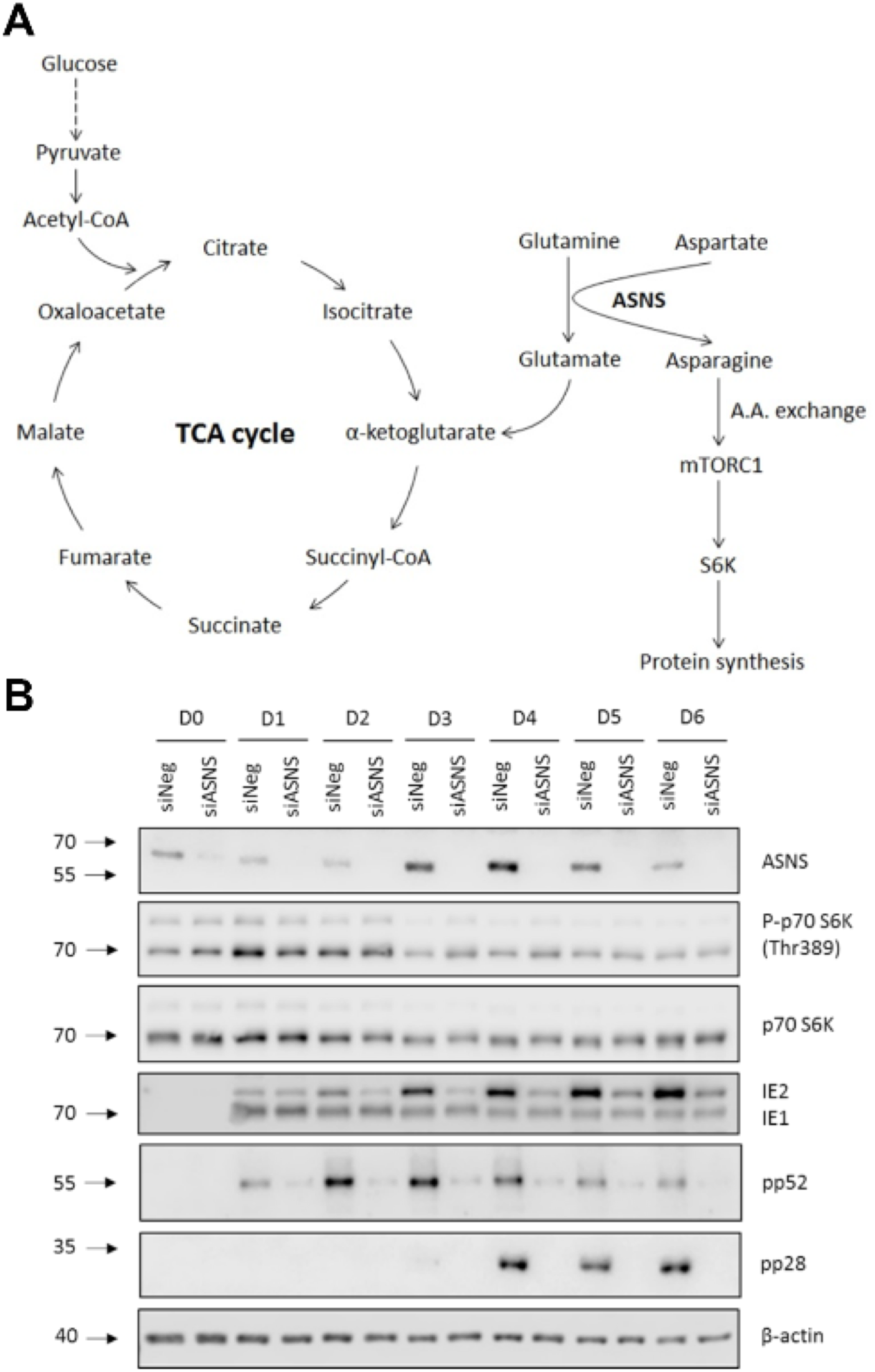
Asparagine depletion did not reduce mTOR signalling during HCMV replication. (A) A schematic diagram of host cell metabolism involving glycolysis, TCA cycle and asparagine biosynthesis. Glucose is metabolised via glycolysis to produce acetyl-CoA which enters the tricarboxylic acid (TCA) cycle (also known as Krebs cycle). Glutamine is metabolised to glutamate, which can be further reduced to make α-ketoglutarate replenishing the TCA cycle. Generation of Asparagine requires conversion of glutamine to glutamate. (B) NHDF cells were transfected with siRNA against ASNS (siASNS) or a scrambled control sequence (siNeg), followed by infection 48 hours post-transfection with HCMV TB40/E-GFP at an MOI of 5. Total protein was harvested at indicated time points. Levels of phosphorylated p70 S6K (P-p70 S6K), p70 S6K, immediate-early (IE1/2), early (pp52) and late (pp28) proteins were determined by western blot analysis. β-actin was used as a loading control.

### Addition of asparagine does not fully rescue glutamine deprivation

Previous studies have shown that infection with HCMV results in increased glutamine metabolism and glutamine starvation results in attenuation of virus replication (7). During infection, increased glutamine metabolism compensates for the diversion of glucose from the TCA cycle. However, glutamine is also required for the *de novo* synthesis of asparagine by ASNS (**Figure 6A**). A recent study demonstrated that addition of asparagine could rescue vaccinia virus replication following glutamine deprivation (21). Therefore, we investigated whether loss of asparagine synthesis contributes to inhibition of HCMV replication following glutamine deprivation. NHDF cells were infected with HCMV in overlay media that contained asparagine (N), glutamine (Q), both or neither. While addition of asparagine resulted in a modest rescue of GFP signal, the result indicates that loss of asparagine synthesis is a minor contributing factor to attenuation of virus replication through glutamine deprivation and loss of precursors for the TCA cycle likely represents the major contributing factor (**Figure 7)**.

**Figure 7.**
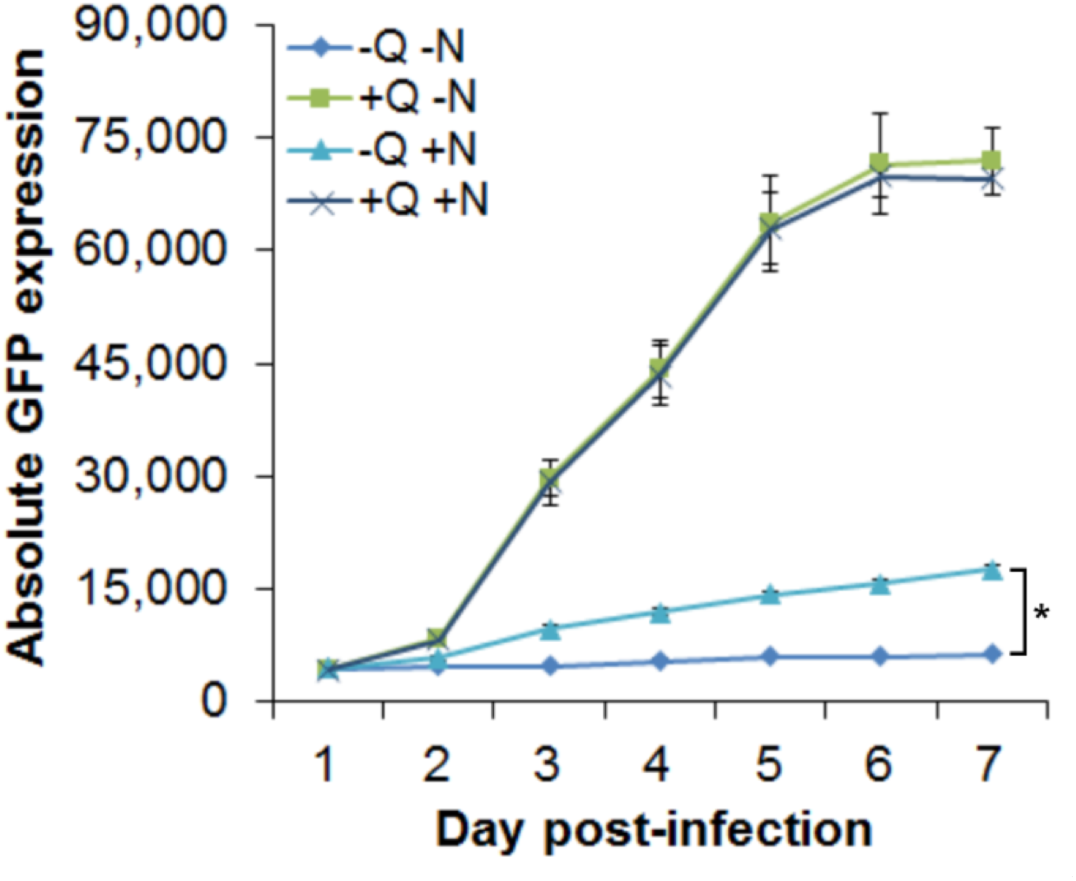
Asparagine supplementation partially rescued HCMV replication in glutamine depleted cells. NHDF cells were grown in media with or without glutamine (Q) and asparagine (N) and infected with HCMV TB40/E-GFP at an MOI of 5 with indicated conditions. GFP fluorescence was monitored every 24 hours for 7 days. *n* = 3; error bars show standard deviations. Two-way ANOVA was used to calculate statistical significance. * = p < 0.05

### Asparagine depletion causes a reversible restriction of HCMV acute replication

Asparagine (Asn, or N) is a non-essential amino acid, meaning it can be produced enzymatically from precursors within cells. The primary fibroblast cells in this study are cultured in media that does not contain asparagine, leaving the cells dependent on this biosynthetic pathway. To determine whether the HCMV phenotype observed following ASNS knockdown was entirely dependent on asparagine deprivation, cells were incubated in media supplemented with 0.1M asparagine two days prior to infection or at the time of infection and maintained throughout the time course. The results show that supplementation with asparagine completely rescued the virus growth phenotype based on GFP reporter expression levels (**Figure 8A**). This result indicates that the phenotype caused by knockdown of ASNS is due to a loss of available asparagine within the cell.

**Figure 8.**
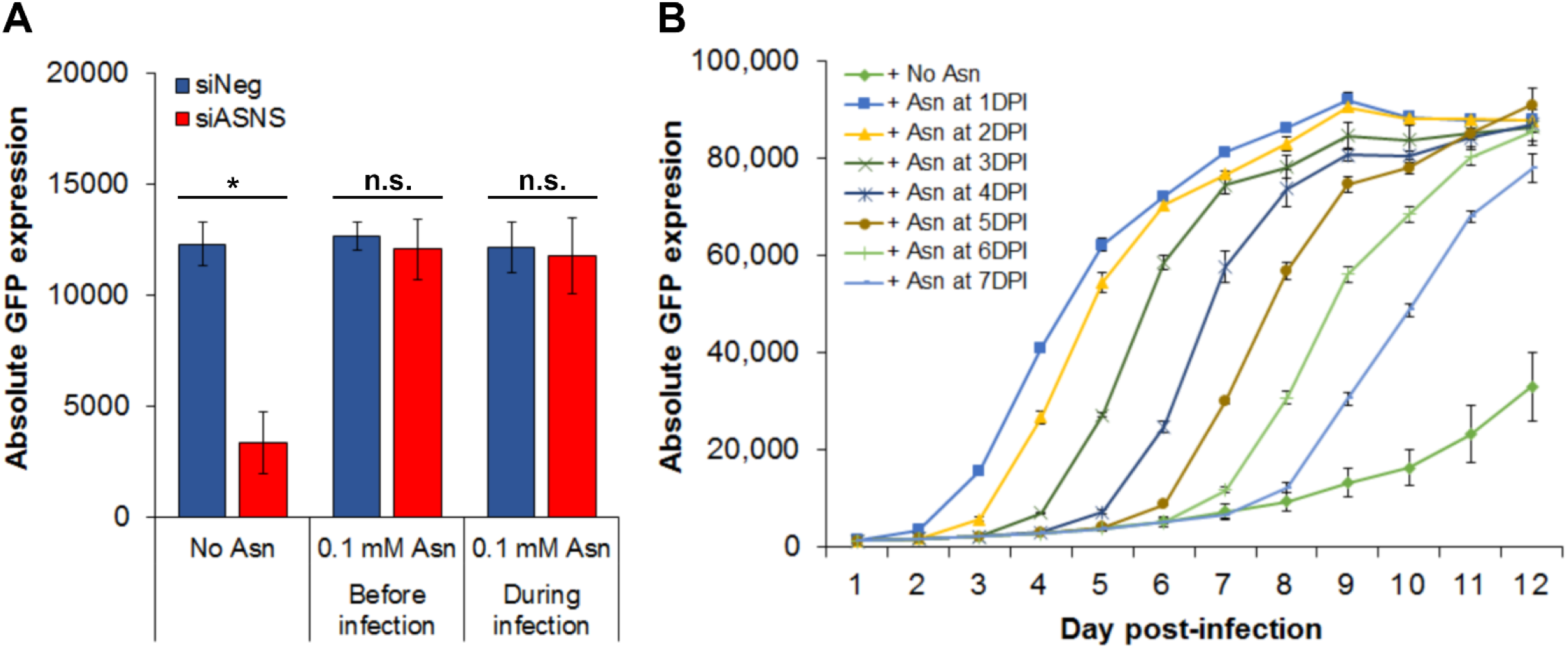
Asparagine deprivation established a reversible inhibition of HCMV infection. NHDF cells were transfected with siRNA pool targeting ASNS (siASNS) or a scrambled negative control siRNA (siNeg), followed by infection 48 hours post-transfection with TB40/E-GFP at an MOI of 5. (A) 0.1 mM of asparagine was supplemented in the medium either at the time of transfection (48H prior to infection) or at the time of infection with primary replication measured at seven days post infection based on GFP expression. (B) 0.1 mM of asparagine was added at indicated time points and GFP fluorescence was monitored every 24 hours for 12 days. *n* = 3; error bars showed standard deviations. n.s. = non-significant. * = *p* < 0.05

As supplementation of asparagine was shown to fully rescue primary replication when added at the time of infection, we wanted to determine whether primary replication could be rescued following extended knockdown of ASNS, when primary replication had stalled for multiple days. Remarkably, full primary replication could be initiated up to seven DPI in ASNS knockdown cells with addition of asparagine, indicating that while virus replication had stalled, infected cells remained viable and the virus was maintained in a state whereby primary replication could be efficiently reinitiated (**Figure 8B**). By seven DPI, primary replication began to recover without the addition of asparagine to the media, likely due to loss of siRNA mediated ASNS knockdown. These results indicate that acute HCMV replication is highly dependent on cellular asparagine levels and deprivation results in inhibition of the replication cycle, prior to DNA amplification and IE2 expression. However, the virus maintains the ability to resume virus replication when asparagine is supplied and reaches peak levels of replication based on GFP reporter gene expression, even following prolonged conditions of asparagine deprivation.

## Discussion

Identification and characterisation of novel host-virus interactions provide valuable insights into how viruses replicate, can inform about the functions of basic cell biology and potentially identify targets for antiviral interventions. Using a high-throughput siRNA screen targeting 6,881 genes, we identified multiple host factors important for HCMV replication, including ASNS, and demonstrate that the non-essential amino acid asparagine is required for HCMV replication at an early stage. Despite ASNS knockdown, global proteins translation levels were maintained Furthermore, given the block in virus replication occurs relatively early in infection and knockdown cells could still fully support replication of HSV-1 and IAV, suggests the loss of replication is not simply due to loss of protein production, but rather indicates asparagine levels feed into a signalling pathway that influences replication of HCMV. While asparagine depletion had little effect on HSV-1 or IAV replication, a recent report demonstrated that, like HCMV, Vaccinia virus (VACV) is also highly dependent on asparagine levels and knockdown of ASNS resulted in attenuation of virus replication (21). Interestingly, despite the similar phenotype, there appears to be differences in the effects of asparagine depletion on the two viruses. For example, addition of asparagine can fully rescue attenuation of VACV following glutamine deprivation, indicating that increased glutamine metabolism in VACV infected cells is necessary for asparagine synthesis, rather than contributing precursors for the TCA cycle.

Furthermore, attenuation appears to be due to a loss of protein production in VACV infection cells. Therefore, while asparagine deprivation has similar effects on both viruses the mechanisms underlying asparagine requirement may be different between the viruses.

There is an increasing appreciation that amino acids are not just building blocks for protein production, but instead participate in many signalling pathways within a cell, affecting protein translation, cell cycle and even apoptosis (22-25). While mTOR is part of a major signalling pathway affected by amino acid levels, our data indicates that knockdown of ASNS in infected cells does not block HCMV activation of mTOR, therefore loss of mTOR signalling does not account for attenuation of virus replication.

Inactive ASNS has been shown to affect cell cycle (25). Infection with HCMV causes cell cycle arrest between G1 and S phase, and studies have shown that virus replication is blocked during other phases of cell cycle due to a failure in immediate early gene expression (26, 27). However, as loss of asparagine synthesis results in a block in cell cycle at the G1 phase, it is unlikely that alteration of cell cycle by ASNS knockdown explains the attenuation of virus replication. Further studies will be required to determine the precise mechanism by which asparagine deprivation results in HCMV attenuation and potential signalling pathways that may be involved.

HCMV infection remains an important clinical issue, both in immunocompromised individuals and during pregnancy, where spread of the virus to the foetus can lead to serious developmental pathologies. There is currently no effective vaccine and available antiviral therapies have significant issues, including side effects and development of resistance. The generation of new treatments against HCMV is therefore required. Very few antivirals have been developed for use against HCMV since the licensing of Ganciclovir and, of these, the same viral genes are often targeted, reducing the effectiveness of these drugs against resistant strains. An alternative strategy for the development of novel antivirals involves targeting of host genes or metabolites required by the virus for successful replication. Development of resistance against drugs that target host genes would be far more complex, as the virus would have to gain mutations that would compensate for the loss of a required cellular factor. In many cases such mutations may not exist. We have shown that reducing available asparagine levels has a profound inhibitory effect on HCMV replication, suggesting this may be a potential strategy for limiting HCMV replication in patients. A recent study demonstrated that reducing asparagine levels in mice through treatment with L-Asparaginase or through dietary restriction, reduced metastatic spread of tumour cells (28). As discussed, tumour cells demonstrate metabolic alterations similar to HCMV infected cells, with increased anaplerosis and dependence on asparagine (7). While cells are able to make de novo asparagine through ASNS, this study suggests that in certain physiological conditions, cells still require free exogenous asparagine, especially cells with a high asparagine dependence. Therefore, reducing available asparagine levels through treatment with L-asparaginase or by dietary restriction may be an effective clinical approach for treating HCMV infection in high risk patients. Temporary dietary restriction would be a particularly attractive approach given the limited likelihood of serious side effects. While this approach would not eliminate the virus, as demonstrated by rescue of virus replication days later with exogenous asparagine, subduing virus replication in combination with other antiviral drugs may still be therapeutically beneficial.

Finally, it will be interesting to determine the potential impact of asparagine levels, and amino acid metabolism in general, on the establishment, maintenance and reactivation of HCMV during latency. ASNS expression levels are relatively low in many tissues in normal conditions. A recent paper reported effects on amino acid metabolism during acute HCMV replication in primary fibroblast cells and it is clear the virus modulates amino acid levels and metabolism, including induction of ASNS as we show here (8). ASNS expression and asparagine levels have been reported to respond to stress signalling and demonstrate tissue specific differences (29). Additional experiments are underway to characterise the reversible block in viral replication during asparagine deprivation, including viral gene expression and characterisation of the viral genome. It will be of interest to determine asparagine levels and levels of amino acids in general during models of HCMV latency and reactivation to determine whether there is any link between regulation of latency and amino acid metabolism.

## Materials and methods

### Cell culture and virus infection

Normal human dermal fibroblast (NHDF, Gibco) cells were maintained in Dulbecco’s modified high glucose medium (DMEM, Sigma) supplemented with 10% foetal bovine serum (FBS, Gibco) and 1X penicillin-streptomycin-glutamine (Gibco). A low passage HCMV strain TB40/E-GFP, which is engineered to constitutively express GFP from an SV40 promoter at the intragenic region between TRS1 and US34, was obtained from F. Goodrum (11) and used for all experiments.

### siRNA screening

The Dharmacon SMARTpool human druggable genome (G-004600-05), cell cycle (G-003250-02) and protein kinase (G-003500-02) siRNA libraries against 6,881 gene targets were prepared in the 96-well format at the concentration of 3 µM (diluted in Thermo Scientific siRNA buffer) at the Division of Infection and Pathway Medicine, University of Edinburgh, followed by the set-up of 384-well master plates at the concentration of 311 nM, using RapidPlate 384 liquid handling robot (QIAGEN). The complete protocol can be found in Virus Host Interactions Methods and Protocols (30). Low passage NHDF cell suspension (approx. passage 11) were reverse-transfected with siRNA and Lipofectamine RNAiMAX (Invitrogen) using MultiDrop 384. At 48 hours post-transfection, media were removed and cells were infected with TB40/E-GFP at an MOI of 5. Relative GFP expression is measured by Cytation 3 cell imaging multi-mode microplate reader (BioTek) every 24 hours for 7 days.

### Cell viability assay

Two cell viability assays were performed at 7 days post-infection (DPI). Media in plates were removed and the CellTiter-Blue reagent with fresh media were added by MultiDrop 384. Following 1-4 hour incubation, the fluorescence at 560/590 nm was measured by Cytation 3 cell microplate reader (BioTek). The relative cell viability was normalised to the reading of control non-targeting siRNA transfected cells.

### Amino acid depletion and supplementation

The DMEM (Gibco; D5796) used for normal cell culture contains a total of 0.876 g/L of L-glutamine and no L-asparagine. An alternative DMEM without L-glutamine (Gibco; D5030) was used to starve cells without L-glutamine and L-asparagine. In preparation of DMEM without L-glutamine, equal amount of sodium bicarbonate (3.7 g/L) and glucose (4.5 g/L) to DMEM (D5030) were added. For L-asparagine supplementation, 0.1 mM of L-asparagine was added, either in DMEM D5030 (to make –Q +N) or in DMEM D5796 (to make +Q +N).

### Western blot analysis

Following transfection, cells were harvested at 0 to 7 DPI in RIPA buffer containing protease and phosphatase inhibitor cocktails (Roche). Protein concentrations were determined by bicinchoninic acid (BCA) assay (Thermo Fisher) following the manufacturer’s protocol. Proteins were separated on 10% SDS-PAGE gels and transferred to nitrocellulose membranes by wet transfer (20% methanol). Membranes were blocked with 5% milk in Tris buffered saline (TBS) and probed with antibodies to HCMV IE1 and IE2 (Merck Millipore; MAB-8131; 1/5000), pp52 (Santa Cruz Biotechnology; sc-56971; 1/1000), pp28 (Santa Cruz Biotechnology; sc-69749; 1/1000), ASNS (Proteintech; 14681-1-AP; 1/1000), p70 S6K (Cell Signalling Technology; 2708; 1/1000), phospho-Thr389 p70 S6K (Cell Signalling Technology; 9234; 1/500), and β-actin (Abcam; ab8227; 1/2500). Secondary antibodies conjugated to horseradish peroxidase (HRP) (Thermo Fisher) or IR800 and IR680 dye (Li-Cor) were used and blots were imaged by Li-Cor Odyssey Fc imaging system. Quantification was done with Li-Cor Image Studio Lite software.

### qPCR

DNA was purified using a DNeasy Blood and Tissue kit (Qiagen) and quantified with a NanoDrop spectrophotometer. The SensiFAST SYBR Hi-ROX kit (Bioline United Kingdom, BIO-92020) and custom gene-specific primer sets were used to assay 20 ng DNA per reaction for HCMV gB (UL55) and GAPDH using the following primers: GAPDH DNA, 5’-GATGACATCAAGAAGGTGGTGA and 5’-CCTGCACTTTTTAAGAGCCAGT; HCMV gB (UL55), 5’-TAGCTACGACGAAACGTCAAAA and 5’-GGTACGGATCTTATTCGCTTTG. Results were normalised to GAPDH DNA levels and then to siNeg levels at 1 DPI by the ΔΔC_T_ method. *n* = 2; error bars represent standard error of the mean.

### STRING analysis

Functional annotation clustering was performed in the free software, Cytospace, with stringApp. The top 115 proviral and antiviral hits were analysed and only interactions with a confidence score of 0.8 or above were shown. To create the network view, each gene was assigned to the cluster with the highest enrichment where the gene was present in the highest fraction of individual clusters. For visualisation purposes, the interactions of a gene with multiple genes in the same cluster (such as the interactions of TAF1 with POLR2A, POLR2B and POLR2G) were removed. Genes that showed no interaction on the network were depicted in diamond shape. The effect of gene knockdown on relative GFP expression of virus replication was indicated in red (reduction) or green (enhancement) on the top right corner of the gene. The functional annotation clusters were arranged in their approximate cellular locations in the final image. Some smaller annotation clusters and unconnected genes (43 genes) were left out due to space limitation.

### Bioinformatics and statistical analysis

siRNA screen data were analysed using Microsoft Excel and its data analytic tools. R studio was used to analyse the correlation between triplicate screens, using the *psych* package. Two-tailed homoscedastic Student’s *t* test was applied to calculate the p-values of the effect of individual gene depletion on HCMV replication. n.s. = *p* > 0.05; * = *p* < 0.05; ** = *p* < 0.005.

## Acknowledgements

This project is funded by MRC and the Principal’s Career Development PhD scholarship by the University of Edinburgh. I’d like to thank Mrs Marie Craigon at the University of Edinburgh for her help with the liquid handling automated machine.

